# Optimal Control of Neural Systems

**DOI:** 10.1101/2022.12.24.521852

**Authors:** Bodo Rueckauer, Marcel van Gerven

**Affiliations:** Department of Artificial Intelligence, Donders Institute for Brain, Cognition and Behaviour, Nijmegen, The Netherlands

## Abstract

Brain-machine interfaces have reached an unprecedented capacity to measure and drive activity in the brain, allowing restoration of impaired sensory, cognitive or motor function. Classical control theory is pushed to its limit when aiming to design control laws that are suitable for large-scale, complex neural systems. This work proposes a scalable, data-driven, unified approach to study brain-machine-environment interaction using established tools from dynamical systems, optimal control theory, and deep learning. To unify the methodology, we define the environment, neural system, and prosthesis in terms of differential equations with learnable parameters, which effectively reduce to recurrent neural networks in the discrete-time case. Drawing on tools from optimal control, we describe three ways to train the system: Direct optimization of an objective function, oracle-based learning, and reinforcement learning. These approaches are adapted to different assumptions about knowledge of system equations, linearity, differentiability, and observability. We apply the proposed framework to train an in-silico neural system to perform tasks in a linear and a nonlinear environment, namely particle stabilization and pole balancing. After training, this model is perturbed to simulate impairment of sensor and motor function. We show how a prosthetic controller can be trained to restore the behavior of the neural system under increasing levels of perturbation. We expect that the proposed framework will enable rapid and flexible synthesis of control algorithms for neural prostheses that reduce the need for in-vivo testing. We further highlight implications for sparse placement of prosthetic sensor and actuator components.

## 1 Introduction

Closed-loop recording and stimulation in neurotechnology is becoming increasingly feasible [44] and calls for algorithmic advances in developing feedback controllers that make best use of improved recording and stimulation capabilities. Studies in human and animal models come with a number of practical and ethical hurdles that motivate the development of algorithms in simulation prior to deployment in a clinical setting. Neural systems, from single neurons to large-scale populations, are characterized by complex dynamics that can be challenging to model and control [43]. Classical control theory [3, 24, 9] provides powerful tools for designing control laws, but they are often limited to linear(-izable) or lower-dimensional systems and require knowledge of the underlying dynamics [47]. The present paper addresses this issue by using scalable, recurrent neural networks (RNNs) as building blocks which are trained via an optimal control oracle or reinforcement learning (RL) [55].

We propose a control-theoretic framework to study the stability and controllability of biologically motivated artificial neural systems embedded in simulated environments. From a high-level perspective, this framework models the brain-machine-environment interaction. We first consider the problem of modelling a neural system to perform a behavioral task in a virtual environment. In the language of control theory, the neural system forms a feedback controller in closed loop with the environment process. In a second step, we simulate a deterioration of the neural system (e.g. at the sensor or actuator) and add a secondary controller (a prosthesis) with the goal to restore behavioral function. In doing so we account for uncertainty in the model of the brain, nonlinearities, measurement noise, and limited availability of observable states and controllable neurons.

Our approach is to develop an in-silico control synthesis framework which builds on tools from optimal control theory and reinforcement learning. The latter is particularly suited because in general we cannot assume knowledge of the environment and brain dynamics, and may only have access to some observations from the environment, the actuator output from the neural system, plus perhaps a temporally sparse reward signal. Even operating under these constraints, RL in principle enables learning arbitrarily complex nonlinear models. In Q-learning [62], a strategy underlying many state-of-the-art RL algorithms, an agent learns a control policy that optimizes a Hamilton-Jacobi-Bellman equation online and without knowing the system dynamics. In optimal control theory, equations of this type are solved analytically (offline and assuming system knowledge) to derive optimal controllers. This common class of equations forms the link between RL and optimal control [31]. Examples for the use of RL in neural control include [41, 54] (in vivo), [65] (in vitro), and [36, 10] (in silico). These studies show great promise of RL to reduce the need for repeated calibration and instead adapt controllers autonomously to changes in the neural code or the sensor / actuator space.

A distinguishing feature of our approach is the unified treatment of environment, brain, and prosthesis as dynamical systems using (stochastic) differential equations. Most of previous work focused either on the brain-environment loop (e.g. to model neuronal dynamics) [48, 46], or on the brain-prosthesis loop (e.g. to train a neural decoder to control a prosthetic limb) [29, 28, 17]. Combining the two loops as proposed here enables end-to-end optimization of the brain model and controller, with adaptation to noise or deterioration appearing in any of the subsystems. The proposed framework provides a substrate for neural control engineering to exploit recent advances in brain-inspired RL [7], safe RL [15, 16], few-shot learning [60], and continual learning [61, 59, 38, 58].

## 2 Methods

### 2.1 Using dynamical systems to model the brain-machine-environment interaction

Before considering the full brain-machine-environment loop, we first concentrate on the unimpaired case and model how an agent (represented by its neural system) interacts with the environment to generate meaningful behavior. From a high-level perspective (Figure 1), the neural system transforms state observations **y**_E_ from the environment into motor commands **u**_E_. Both the neural system and the environment can be modeled by controlled stochastic differential equations (SDEs) of the form

**Figure 1:**
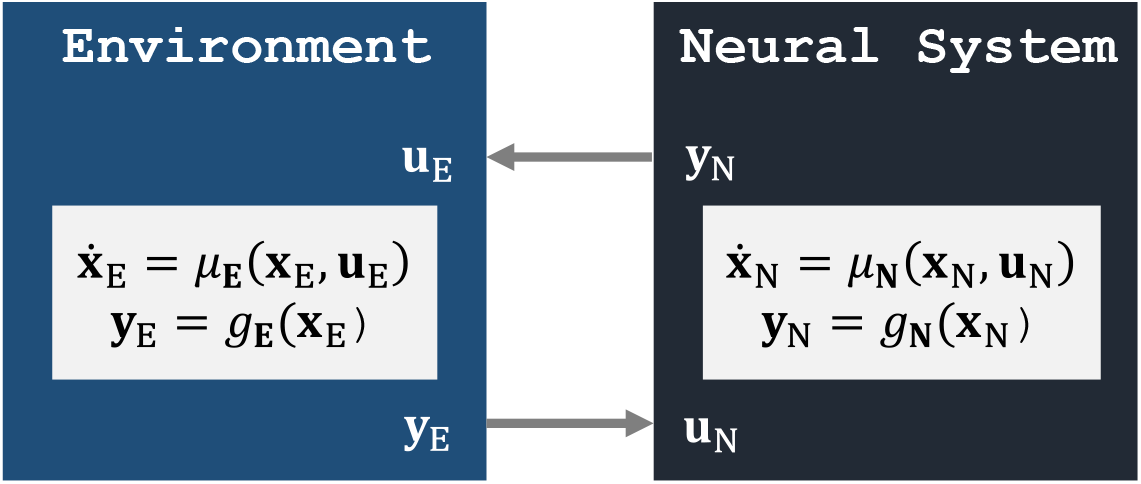
Classical agent-environment loop, where a neural system interacts with its environment by observing its states and applying feedback control. Here, the deterministic case is shown for simplicity; see Equation (1) for the stochastic form.

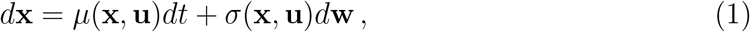

where the drift *μ* models the evolution of the states **x** under some external input **u** and the diffusion *σ* models how Brownian noise **w** enters the system. In the absence of noise, the model reduces to the ordinary differential equation 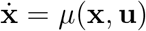.

So far, we formulated the system dynamics as *continuous* differential equations. Various discretization methods can be applied for the purpose of simulating the system dynamics on digital hardware or solving for the states **x** numerically. A common approximation for SDEs is afforded by the Euler-Maruyama method. Starting from some initial value **x**_0_, the solution is recursively defined as

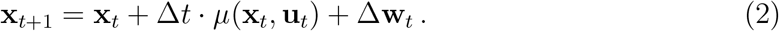

Here, Δ*t* = *T/N* is the step size obtained by dividing the integration interval *T* into *N* steps. The noise term Δ**w**_*t*_ is drawn from a normal distribution 𝒩(**0**, Δ*t S*) with 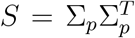. Here we assume a fixed diffusion matrix *σ*(**x, u**) = σ_*p*_ with σ_*p*_ the process noise covariance.

Closed-loop control of a dynamical system requires observing its states. In real-world scenarios, one typically has access to just a subset of the states, which may moreover be perturbed by sensory noise. The resulting observations can be described as

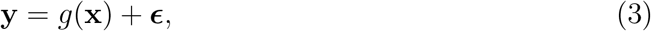

where *g*(**x**) could be a matrix-vector product *C***x** and *ϵ* ∼ 𝒩(**0**, σ_*o*_) with σ_*o*_ the observation noise covariance.

### 2.2 Example environments

To illustrate the general applicability of the concepts developed in this paper, we introduce one linear and one nonlinear example of dynamical systems, which will be used to represent an environment for the neural system to act in.

#### 2.2.1 Double integrator for particle control

A canonical example from control theory is the double integrator, which models the dynamics of a particle in a one-dimensional space under the influence of an external force **u**. The environment’s states are given by the particle’s position *q* and velocity 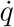. Their dynamics are determined by the differential equation 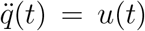, which identifies the particle acceleration with the control force. For the stochastic double integrator, the combined state 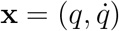 evolves according to

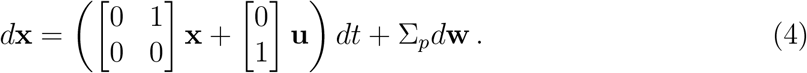

Given a moving particle away from the origin (**x**(0) ≠ 0), the task of the neural system consists in determining the sequence of motor commands **u**(*t*) that force the particle to the origin and stabilize it there (**x**(0) = 0). The task can be made more challenging by increasing the process noise covariance and by allowing the neural system to observe only a noisy version of the position but not the velocity:

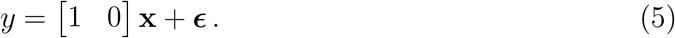

We will discuss ways to solve this task in Section 2.4 below.

#### 2.2.2 Balancing an inverted pendulum

Another classic benchmark for control algorithms is the inverted pendulum, or cartpole balancing problem [4]. It consists of a cart with a pole attached that has its center of mass above the pivot point. The task is to move the cart horizontally so as to keep the pole in its unstable vertical position. When limiting the pendulum to one degree of freedom, the system has four states: cart position *x*, velocity 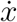, pole angle *θ*, and angular velocity 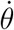. The system is nonlinear: In absence of control forces, the angle evolves as 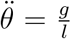 with standard gravity *g* and pole length *l*. Thus,

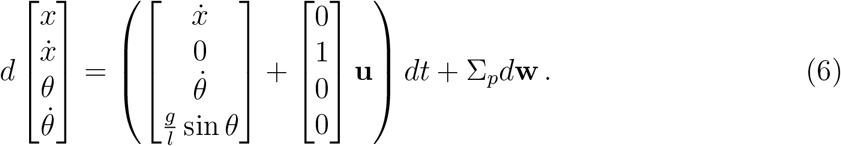

### 2.3 Analytic solution of linear, known systems

For a linear system with known dynamics, an optimal direct feedback controller can be derived analytically. The central idea is that the controller should minimize the squared deviation of states from their desired values, while expending the least amount of energy to do so. In the case of static target states, the problem is called Linear Quadratic Regulator (LQR), with the quadratic cost function

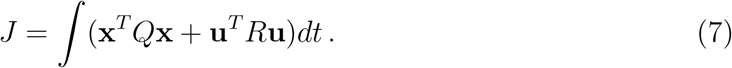

The matrices *Q, R* weigh the contribution of individual states and controls to the overall cost. In general, adjusting these parameters to a particular problem can be challenging; for the particle control we find a uniform weighting *Q* = *R* = *I* sufficient. The LQR cost function is minimized by a simple proportional feedback rule **u** = *K***x**. The LQR gain matrix *K* can be computed using standard numeric libraries.

In Equation (7) we assumed full access to the states, i.e., *g*(**x**) = **x** and ***E*** = **0** in Equation (3). More realistic is the case of partial noisy observations. Fortunately, the optimal solution may still be applied by combining the controller with a Kalman filter, which enables estimation of unobserved and denoising of observed states in linear systems. Given the system dynamics, the Kalman filter operation can be applied via a simple matrix-vector multiplication 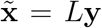, mapping noisy partial observations to full state estimates, which can be used in the LQR loss (7). The combination of LQR with a Kalman filter is called Linear Quadratic Gaussian (LQG). In the next Section we consider the case where linearity and system knowledge cannot be assumed.

### 2.4 Modelling the neural system using a recurrent neural network

Research in control theory over the past decades has resulted in a wealth of techniques to solve tasks like the ones described in Section 2.2. Which approach to take depends on the stringency of assumptions we make about the system. At one end of the spectrum, one can compute an optimal control policy analytically if the system is linear and the state equations are known (Section 2.3). The controller (or neural system in our terminology) is then a simple matrix that maps the state observation vector **x** to a control signal **u**. At the other end of the spectrum, one can employ reinforcement learning to find a control policy for a nonlinear system with unknown dynamics and partial noisy observations. In that case, the neural system controller is usually an artificial neural network with a large number of parameters.

Here, we use recurrent neural networks (RNNs) to model the controller. RNNs have a long history in modeling of neural systems [52] and use in control applications [35, 53], making them an ideal candidate in the context of this paper. We will see in Section 2.5 that these networks are compatible with both ends of the spectrum - they can adapt to analytic optimal control laws just as to data-driven policy search through RL. In particular, the statefulness of RNNs turns out to be a crucial component in dealing with partially observed and/or noisy observations.

The continuous-time RNN used here follows work by [52] and is defined by the ordinary differential equation

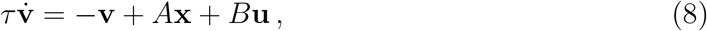

where the vector **v** represents the neurons’ membrane potentials, **x** = *h*(**v**) are the firing rates obtained by passing the membrane potential through a sigmoidal activation function *h*, and *τ* is the time constant of charge integration and decay. The neurons integrate inputs **u** via synaptic weights *B*, and are recurrently connected through the matrix *A*. Layers of such RNN units may be stacked hierarchically, so that the rates **x** of one layer become the input **u** to the next. The network is further assumed to include a fully-connected readout layer

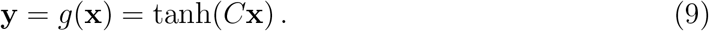

Using a (split-step) Euler method, the continuous-time equation (8) can be discretized. By choosing a step size Δ*t* that matches the time constant *τ*, we obtain an expression for the state evolution that is equivalent to the RNN model commonly used by the AI community:

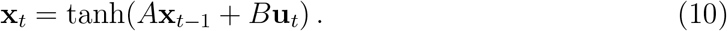

Choosing an RNN as controller has the important consequence that our neural system now becomes a dynamical system itself, just as the environment which it aims to control. This way we can unify the proposed methodology for studying the brain-environment interaction.

In terms of free parameters, the model described by equations (9) and (10) is characterized by the synaptic weight matrices *A* (recurrence), *B* (input), and *C* (output). In the context of this paper, we find it useful to interpret the input component *B***u** as sensory subsystem, the output layer *C***x** as motor system, and the recurrent term *A***x** as an association area. Figure 2 illustrates these RNN components within the original control loop of Figure 1. The next Section describes three approaches to fit the model parameters **θ** = {*A, B, C*}. Note that biases are included in the models but not shown in the equations to simplify notation.

**Figure 2:**
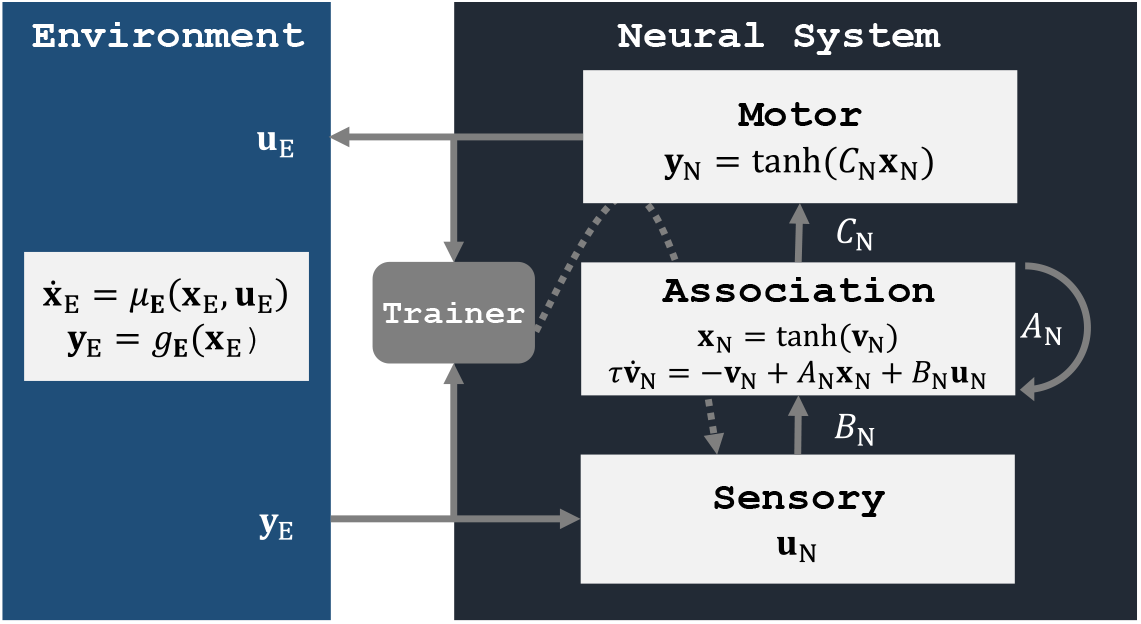
Both the environment and the neural system are modelled as dynamical systems using ordinary or stochastic differential equations. Here, the neural system is represented by an RNN with a sensory input component, a motor output component, and an association area which provides the input-output mapping. The system can be trained using backpropagation through time [63].

### 2.5 Learning a control policy in the neural system

While control theory provides tools to synthesize controllers for systems that are nonlinear, only partially observable, and whose dynamics are unknown, these methods often include an attempt to linearize the system around an operating point, perform system identification, or include an (extended / unscented) Kalman filter [23] to estimate unobserved states. If successful, these approaches result in a system where the control policy can be derived analytically, and often provide performance guarantees and safety / stability regions. On the other hand, they may not always be applicable or scale well to high-dimensional systems. The considerable complexity of the multi-stage processing pipeline makes some of these approaches difficult to maintain and to transfer between problem domains. Training an RNN controller using RL has the benefit of being conceptually simple and at the same time highly scalable. It assumes no linearity or prior knowledge of the system dynamics. Due to the persistent states of the neural network model, the RNN controller performs temporal integration and can adopt the role of a Kalman filter, estimating unobserved states while filtering out noise in the observed ones.

The present paper aims to study how a neural system (the primary controller) can be stabilized by a secondary controller in the presence of perturbations. In this context we do not attempt to find the best tool from classic control theory to solve the primary control problem. We focus instead on obtaining a neural-network-based controller that can serve as test bed for developing the secondary controller. These considerations motivate the use of an RNN-based controller.

Once settled on an RNN as neural system, the training method is usually going to involve a form of stochastic gradient descent with back-propagation through time [63], which updates the network parameters to optimize some objective function. This generic trainer component is depicted in Figure 2, with the gradient flow marked by a dashed arrow. The question that remains is how motor commands **u**_E_ and environment observations **y**_E_ are combined to obtain a learning signal. To answer it we need to consider the properties of the environment on which the controller acts, and how much knowledge of the environment dynamics we can assume. These considerations, together with the possible training methods, are listed in Table 1 and described in more detail in the following sections.

**Table 1:**
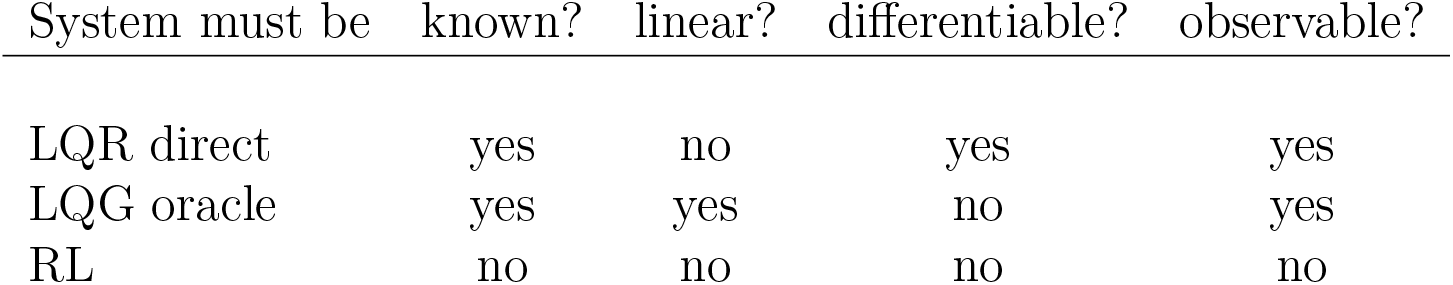
Selecting a training method for the RNN neural system depends on properties of the environment. Note that observability can be achieved by adding a (non-)linear Kalman filter, provided that the system equations are known.

| System must be | known? | linear? | differentiable? | observable? |
| --- | --- | --- | --- | --- |
| LQR direct | yes | no | yes | yes |
| LQG oracle | yes | yes | no | yes |
| RL | no | no | no | no |

#### 2.5.1 Direct optimization of LQR cost

We consider first the most optimistic case, where the state equations (1) are known and we assume that we may differentiate through the environment. Because the neural system is a differentiable RNN, the entire brain-environment model becomes end-to-end differentiable, allowing direct optimization of the parameters using automatic differentiation [5]. That is, after defining a suitable loss function, the neural system can be trained via gradient descent, with the environment in the loop. The samples for training can be generated on the fly: As the neural system steps through the environment, it observes states **x**_*t*_ and produces motor commands **u**_*t*_, which are passed to the loss function to generate a learning signal that is propagated back through the network. In the backwards pass, only the parameters in the neural system are updated; the environment parameters are frozen.

We now turn to defining the objective function. For a linear regularization problem like the particle stabilization described in Section 2.2.1, we can simply choose the LQR cost (7). The hypothesis is that the neural system will learn a control policy that follows the analytically derived optimal solution.

In case of partial or noisy observations, the same objective function may be used. We can estimate the latent state explicitly by including a Kalman filter, whose matrix-vector-product operation is straightforward to implement in a neural network and thus integrates well with the direct optimization approach. Alternatively, a Kalman filter can be learned implicitly by the stateful RNN as we demonstrate in Section 3.1.2.

#### 2.5.2 LQG oracle

Next we consider the case that direct optimization on some loss function is not possible because the environment is not differentiable, for instance due to discontinuities in the dynamics. We may still be able to exploit tools from optimal control theory like the LQR and Kalman filter introduced in Section 2.3. As before, we assume knowledge of the state equations (1). In addition, the system should either be linear or linearizable. Full observability is not a requirement as a Kalman filter is available under the current assumptions.

Following Section 2.3, the optimal LQG controller, could be used directly as surrogate neural system. Here, however, we illustrate the case of using the analytically derived optimal controller as teacher for the RNN-based neural system.^1^ The LQG teacher can be deployed in the environment to collect a dataset of inputs **y** and labels **u**, which are then used for conventional supervised training of the RNN.

#### 2.5.3 Reinforcement learning

The most general case, requiring the fewest assumptions, is to use RL for training the neural system RNN. Reinforcement learning is a form of machine learning which only requires reward signals *r*_*t*_ as typically encountered by agents in a realistic environment, rather than labeled input/output pairs as required in supervised learning. An agent in state *s* learns to take actions *a* according to a policy *π*(*a, s*) = Pr(*a*_*t*_ = *a*|*s*_*t*_ = *s*) which maximizes the discounted future reward 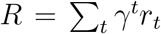. It does so by striking a balance between exploring new states and actions, and exploiting those already known to have high value. The optimization of a value function *V*_*π*_(*s*) = 𝔼[*R* | *s*_0_ = *s*] links RL to the field of optimal control and combines it with the benefits and drawbacks of data-driven model-free control.

In terms of the criteria in Table 1, RL is applicable to linear or nonlinear systems with known or unknown dynamics. While there are no guarantees for the learning performance, a noisy partial observation of states together with a scalar reward signal after possibly long time intervals is usually enough to train the neural system.

In the present work, we employ the proximal policy optimization (PPO) algorithm [49], which is an established online policy-gradient RL method that is relatively easy to use while achieving state-of-the-art results. Other benefits of PPO include its support of both discrete and continuous action spaces, and compatibility with recurrent models. We adapted the Stable-Baselines3 [42] implementation of PPO to work with our RNN model.

### 2.6 Restoring behavior of a perturbed neural system using a secondary controller

The previous sections outlined various approaches of training a neural system to perform meaningful behavior in an environment. With this model at hand we can now study ways to maintain the system performance when parts of the neural system begin to degrade.

Impairment of a biological sensory-motor controller can take many forms, such as cell death or loss of sensor function. Plasticity and a drift in neural representations call for controllers that coadapt with the neural code [51]. Here we implement model perturbation in a generic way by adding univariate Gaussian white noise of various strengths to the synaptic weights of the trained neural system. We consider perturbation of the association (*A*), sensory (*B*) and motor (*C*) populations separately.

Perturbing the neural system can be seen from a control-theoretic perspective as increasing model uncertainty in the SDE (1). The brain-environment loop (Figure 2) is then described by a set of coupled SDEs and can be redefined as a new composite dynamical system. With this unified view we can apply the same control techniques as earlier for the brain-environment interaction, but now with the aim to restore the function of the impaired neural system. Specifically, we add a secondary RNN controller as shown in Figure 3. It follows the same state equations (10) as the neural system RNN, but its components can be interpreted in terms of a neural prosthesis. The input **u**_P_ takes on the role of a neural recording device such as microelectrodes or neural imagers. The output component **y**_P_ represents a stimulation device. The hidden RNN layer closes the loop between neural recording and stimulus generation.

**Figure 3:**
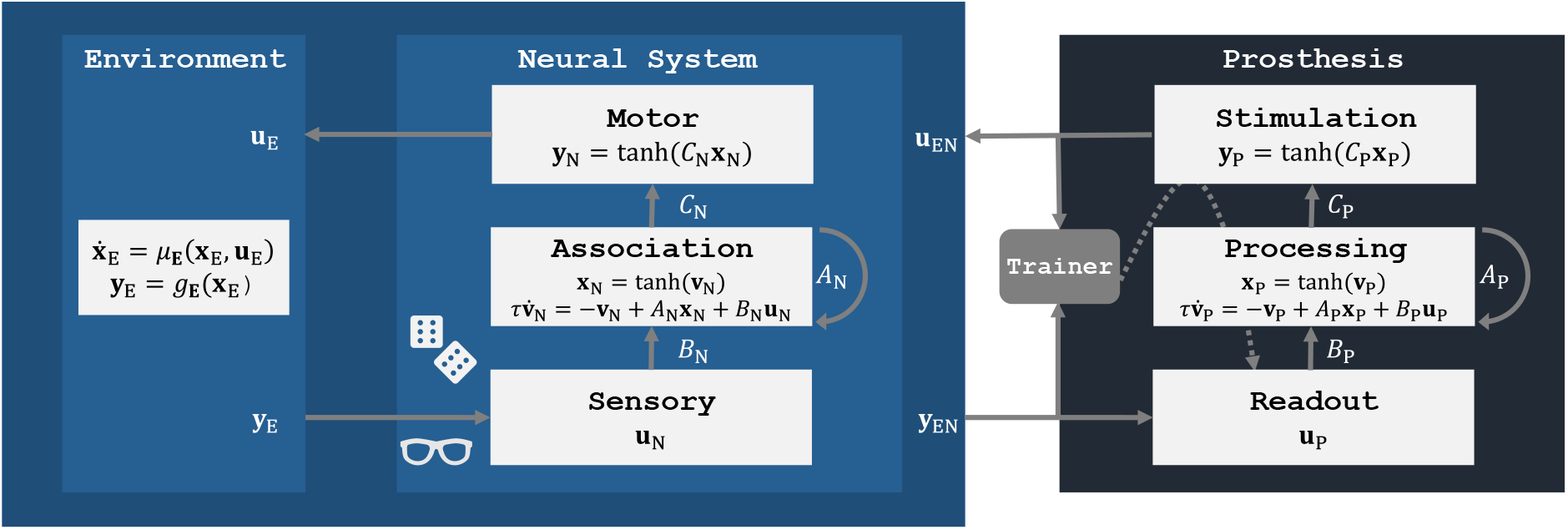
Framework to model a perturbed neural system in interaction with its environment (left) and restoring its performance by training a secondary controller (right).

The observations **y**_EN_ of the composite brain-environment system could be a combination of states from both subsystems. For instance, a visual prosthesis would process a camera feed of the environment in addition to neural activity measurements. In our experiments we read out the states **x**_N_ from the association layer:

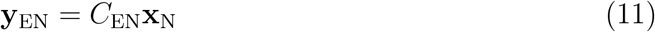

with *C*_EN_ = *I*, assuming full observability initially. We show in Section 3.3 that equivalent performance is achieved when only a small fraction of neurons can be observed and controlled.

Another design choice concerns the feedback control signal **u**_EN_. Following our interpretation of neural stimulation, we apply it in form of charge injected in neurons of the association layer:

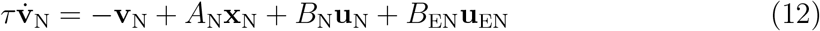

with *B*_EN_ = *I*. We use the same training approach for the prosthesis as for the neural system, i.e., if the latter was trained using RL, the former is as well. The neural system parameters remain fixed after perturbation, only the parameters of the secondary controller are allowed to evolve.

## 3 Results

In this Section we outline first how the neural system was trained on the two tasks described in Section 2.2. The brain model is then perturbed and a prosthetic controller is trained to restore the function of the neural system. Finally, we consider the case of an underactuated system, where the number of controls is lower than the number of neurons. Code to reproduce the results shown here is available online^2^.

### 3.1 Training the neural system

#### 3.1.1 Direct optimization of LQR cost

For stabilizing a drifting particle at the origin, we use a neural system consisting of 50 artificial neurons, which form the association layer in Figure 2, and follow the dynamics (10). The sensory signal **y**_E_ here consists of the full state **x**_E_, i.e., position and velocity of the particle. The motor system is represented by a single neuron which receives input from the association population. Following Equation (9), this neuron produces the real-valued control signal **y**_N_ = **u**_E_ representing the acceleration applied to the particle. The environment (4) is implemented in Python, using the Euler method to approximate the continuous dynamics. The RNN neural system was trained for 10 epochs with the Adam optimizer. During each epoch, the system traversed 10k trajectories in state space, with initial starting points sampled at random from a grid of 100 *×* 100 locations.

The training result is shown in the first row of Figure 4. The panels on the left show example trajectories of the drifting particle in phase space. When the neural system has not been trained (dashed line), the particle drifts off with uniform or increasing speed along the x-axis. After training directly on the LQR cost (solid line), the particle is reliably brought to rest at the origin (marked by a small cross). The trajectory then closely matches the analytically derived optimal control baseline (dotted line).

**Figure 4:**
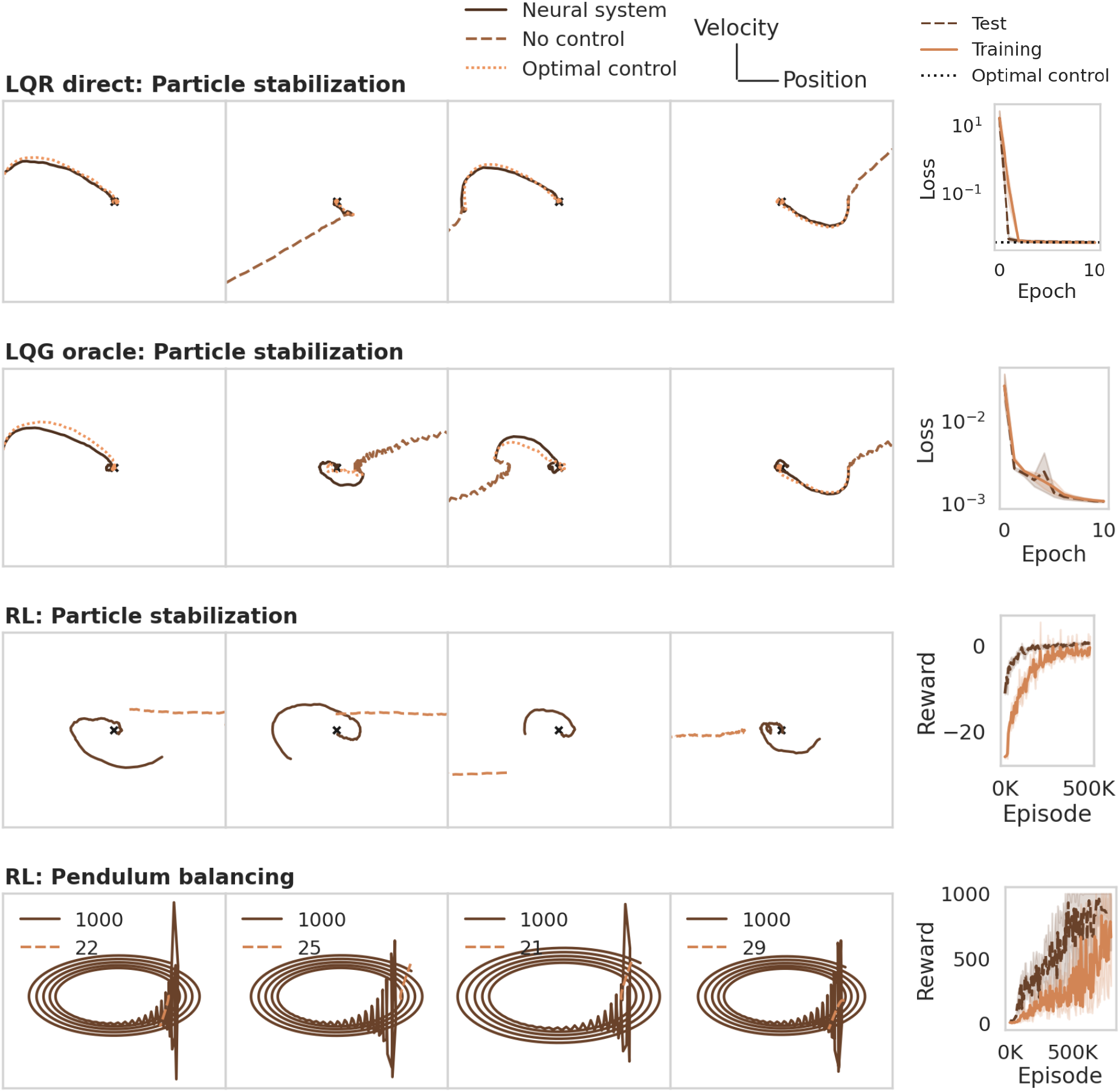
Learning a control policy in the neural system. The panels on the left illustrate the evolution of the environment in state space. The panels on the right show the training curves of the RNN neural system. The rows illustrate the training methods discussed in Section 2.5, applied to the particle stabilization and pendulum balancing problem. The last row also shows the final rewards. In all four cases, the RNN system succeeds in learning a control policy to solve the task.

#### 3.1.2 LQG oracle

To illustrate the oracle approach, we trained an RNN controller with the same specifications as in Section 3.1.1, with three main differences. First, only the position but not the velocity of the particle was observed, and overlaid with Gaussian white noise of variance 0.1, as in Equation (5). Second, rather than training online, the network was trained on batches of pre-collected data from an LQG optimal controller. Third, rather than using the LQR cost directly, we computed the mean squared difference between the control signal generated by the neural system and the control signal from the LQG oracle.

The second row in Figure 4 illustrates the training result. In this case of partial noisy observability, the neural system again succeeds in stabilizing the particle while following the optimal control baseline closely.

#### 3.1.3 Reinforcement learning

For the RL approach we use the same architecture as in the previous sections. Again, the sensory signal is limited to a noisy version of the particle position. The neural system is trained for 500k episodes with a linearly decaying learning rate of initially 2 · 10^−4^. All other hyperparameters of the PPO algorithm were left at their default value.

An important component of training RL agents is to design the reward function. Here, the reward is given by the negative LQR loss, plus a reward of 1 when stabilizing within a distance of 10^−3^ from the origin, at which time the episode terminates as successful. If the particle fails to stabilize within 100 steps, the episode times out and is considered a failure. Row three in Figure 4 illustrates that the sparse RL rewards are sufficient for the RNN to learn the regularization task.

Unlike the particle stabilization problem, the inverted pendulum (cf. Section 2.2.2) is nonlinear. To solve this second task, the only changes we make to the RL training pipeline are to increase the size of the association population to two layers of 128 recurrent neurons each and increase the training duration to 800k episodes. The sensory signal **y**_E_ consists of the position *x* of the cart and angle *θ* of the pole, but not their respective velocities. The association population again projects to only one motor neuron, which produces a real-valued force to steer the cart left and right. To simulate the inverted pendulum environment, we use the MuJoCo [57] implementation within the OpenAI Gym interface^3^. The reward is defined as the number of steps for which the pole can be kept upright. A balancing episode is aborted as unsuccessful when the pole angle *θ* exceeds 11°. Successful episodes time out after 1000 steps, achieving the maximum reward of 1000.

We observe in row four of Figure 4 that the pole tips over within few steps (*r* < 30) when controlled by a randomly initialized RNN. The trained neural system is able to stabilize the pole up to the time limit of 1000 steps. In phase space, a successful trial is often characterized by a harmonic oscillation, and a failure by an outward spiral or straight line.

### 3.2 Training the prosthetic controller

The result of training the prosthesis directly on the LQR cost after perturbing sensor, association, and motor populations is shown in Figure 5 for the particle control task. The panels on the left show example trajectories for increasingly strong perturbation levels. The final column compares the performance of the perturbed system with and without a prosthesis against the optimal LQR baseline. The secondary RNN controller is able to restore the behavior of the neural system across a wide range of perturbation levels, regardless where the perturbation was applied.

**Figure 5:**
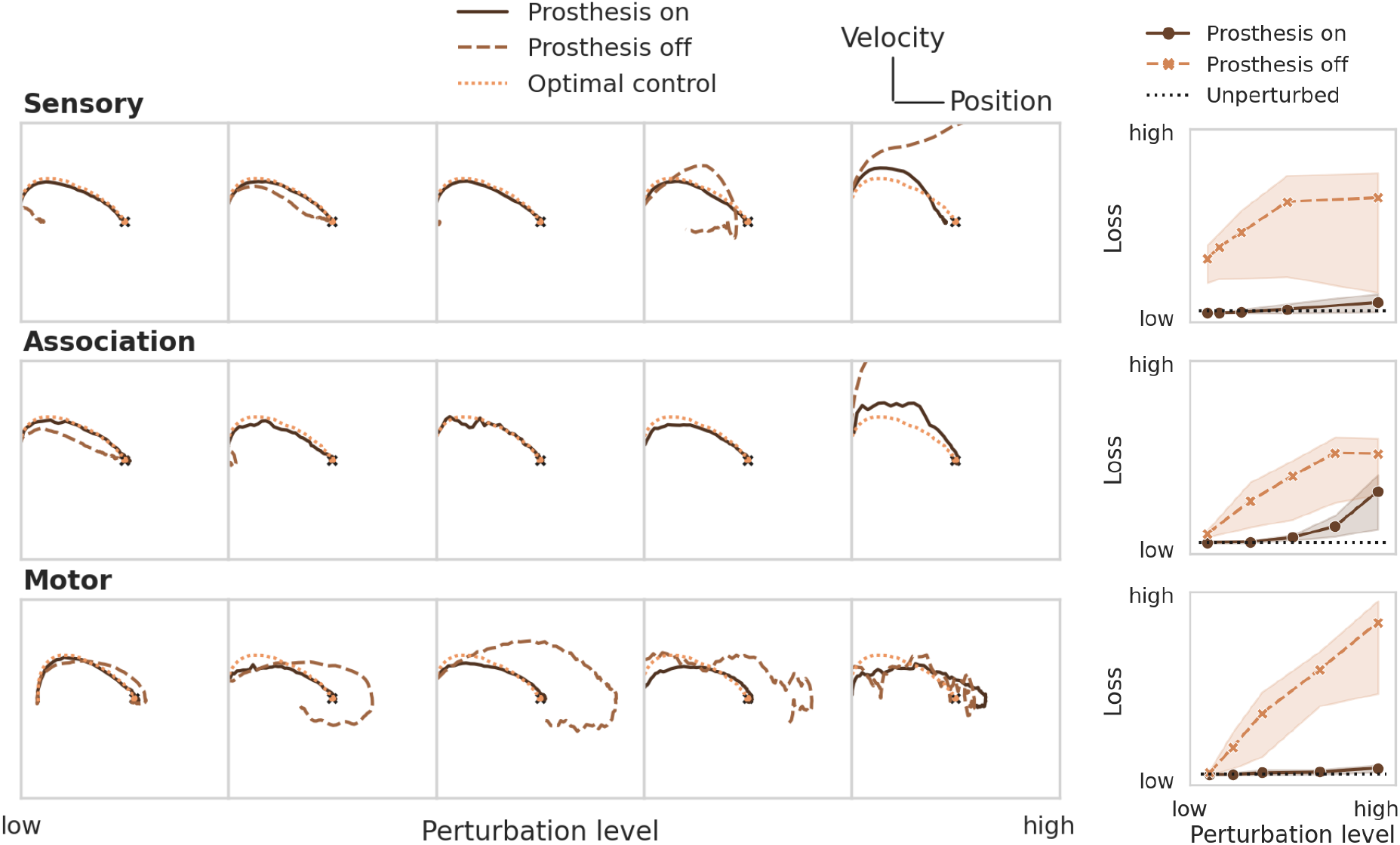
Restoring particle stabilization performance by training a prosthetic controller directly on the LQR cost. The panels on the left show example trajectories at increasing perturbation strengths (from left to right). The rightmost column compares the loss of the system after training the prosthetic controller against the uncontrolled and unperturbed baselines. The three rows illustrate the case of an impaired sensory, association, or motor population.

This direct learning shows what can be achieved under optimistic assumptions (Table 1). In a real-world behavioral setting, the environment dynamics will likely be unknown, nonlinear, noisy, and only partially observable. Then, one can approximate the optimal control solution via reinforcement learning, using only sparse reward signals to learn from. Figure 6 demonstrates that even under these conditions, the prosthesis manages to restore neural system performance. RL training results on the nonlinear pole balancing task are shown in Figure 7.

**Figure 6:**
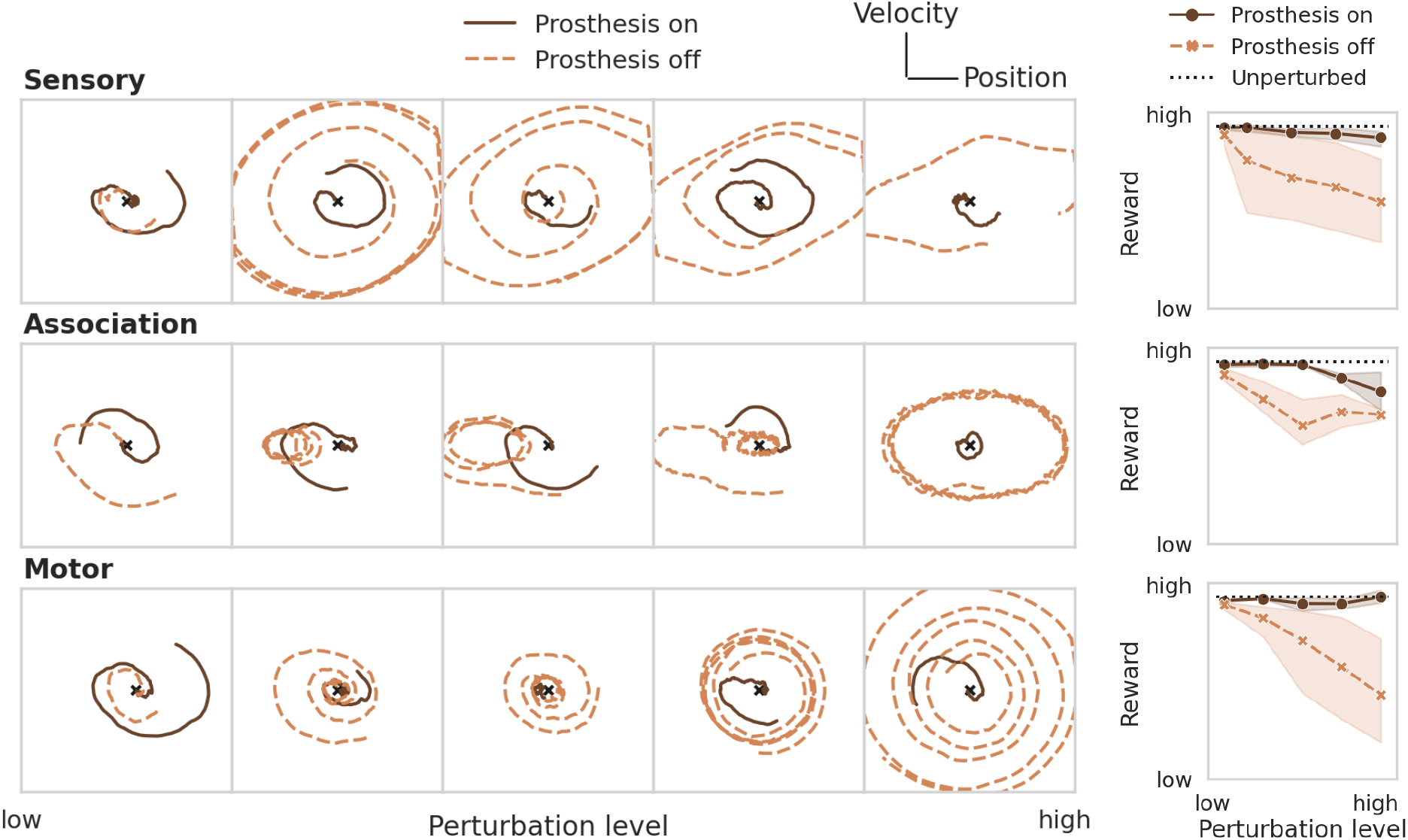
Restoring particle stabilization performance by training a prosthetic controller using RL. See caption of Figure 5 for a more detailed description.

**Figure 7:**
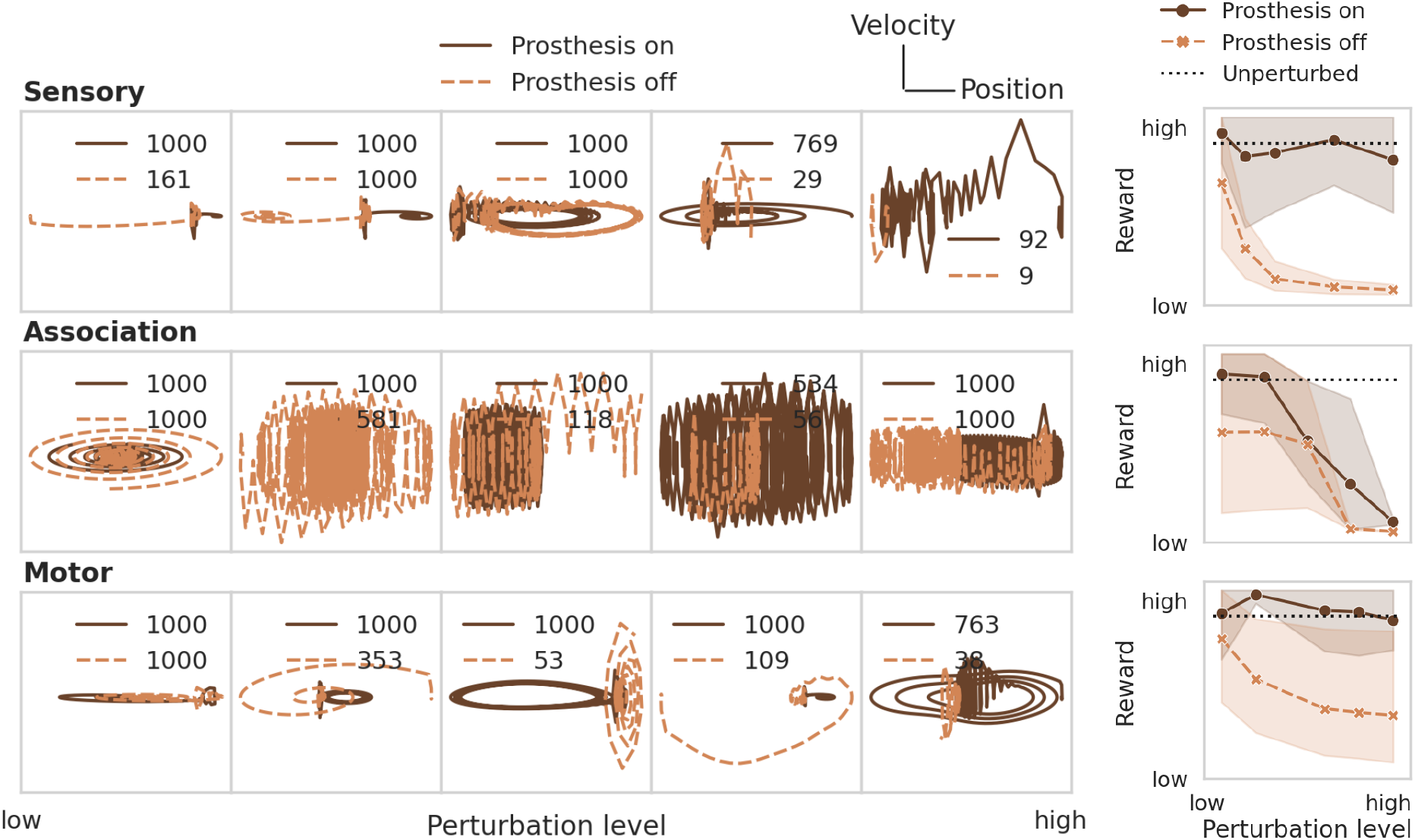
Restoring performance of the neural system on the pole balancing task by training a prosthetic controller using RL. See caption of Figure 5 for a more detailed description.

### 3.3 Underactuated case: Reducing controllability and observability

In the previous section we assumed access to all neurons in the association layer for recording and stimulation. With a microelectrode array for instance, only a limited number of recording and stimulation locations may be accessible. Due to reactions of the neural tissue, the usable set of electrodes may change over time, as well as the set of neurons corresponding to a given electrode. Safety limits for local charge density may add further constraints. See [13] for a review. To account for these properties we reduce the number of observable and controllable states in the association population by removing columns in *B*_EN_ and rows in *C*_EN_. The prosthetic controller is then trained as before with the aim to restore the function of the perturbed neural system. Figure 8 shows the performance after training with different degrees of stimulation and recording electrode coverage. The combined brain-prosthesis system maintains the performance of the unperturbed neural system with as little as 10% of the association neurons being observed and controlled.

**Figure 8:**
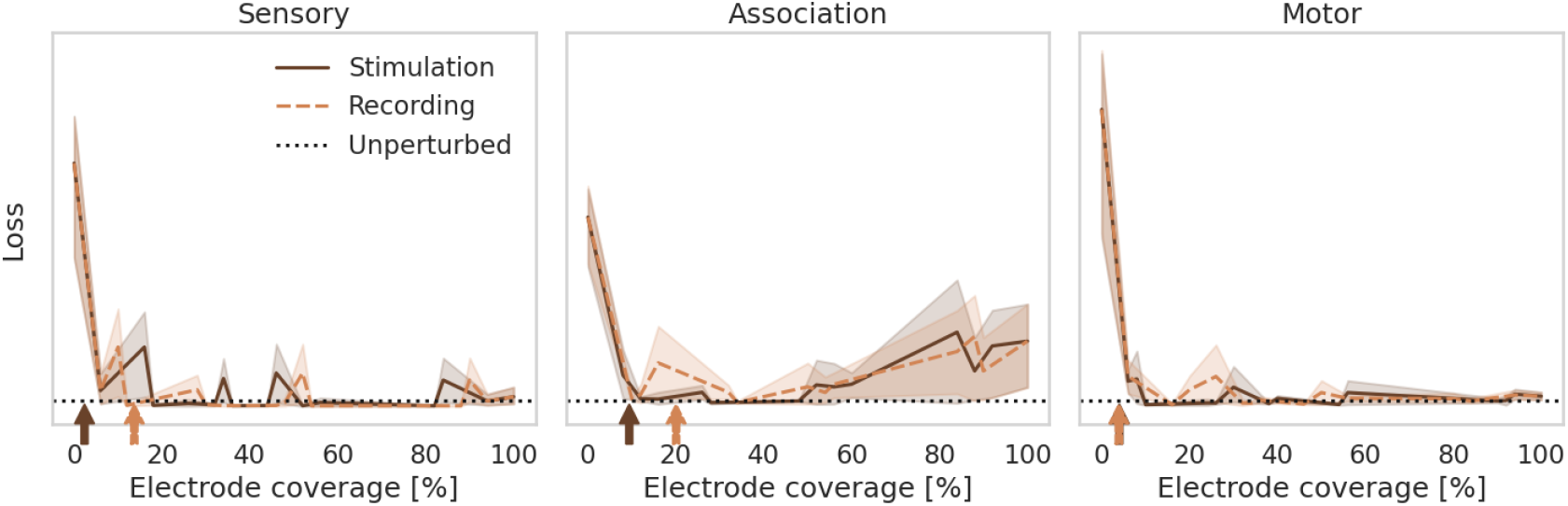
Effect of reducing controllability and observability. The loss is shown when the prosthesis can record from and stimulate only a fraction of neurons in the neural system. The prosthesis restores the performance of the unperturbed neural system when about 10% of the neurons are accessible. This number approximately matches the lower bound estimated from the controllability and observability Gramians (indicated by arrows).

Control theory provides tools to measure the observability and controllability of a controlled system. In a known linear system, one can use the system matrices *A, B, C* to analytically compute an observability matrix 𝒪 and controllability matrix 𝒞 (see e.g. [9]). The system is called observable or controllable, if the corresponding matrix is full rank. In a nonlinear or unknown system, it is possible to estimate an *empirical* Gramian matrix by measuring the system response to stereotypical control impulses [30, 19]. Eigenspectrum analysis of the controllability and observability Gramians reveal directions (given by the eigenvectors with the largest eigenvalues) along which a system can be steered with the least amount of control energy, and observed with the highest signal-to-noise ratio. These directions lie in eigen- rather than state-space, so we cannot use them directly e.g. to select the most suitable subset of electrodes for recording and stimulation. However, the eigenspectrum does provide an indicator for the intrinsic dimensionality of the system and thus the number of electrodes required. Specifically, we can count how many eigenvectors are needed to explain 90% of the variance of the controllability and observability Gramians. This number of dimensions gives a lower bound on the number of electrodes needed for stimulation and recording. We indicate it with an arrow in Figure 8. It turns out that for the particle stabilization problem, this lower bound coincides with the minimum number of electrodes found empirically. In other words, the RNN controller learns to make optimal use of the small number of probes it has available.

## 4 Discussion

The present work proposes a framework based on dynamical systems to model interactions between the brain, its environment, and a prosthesis. Established methods from optimal control theory and reinforcement learning are applied to train the neural system to perform a task and then train a prosthesis to restore its function when the neural system is perturbed. Here we discuss the design choices of this conceptually simple framework as well as implications for the development of neural controllers.

### 4.1 Modelling the neural system and prosthesis

In this work we used an RNN to model both neural system and prosthetic controller. Choosing this class of model has several advantages. An RNN can be described by a set of differential equations, which makes the combined environment-brain-prosthesis system uniform in structure and amenable to a common methodology. The statefulness of RNN neurons facilitates working with temporal data, which is ubiquitous in realistic settings. Statefulness has the additional advantage of enabling the RNN to adopt functions of a Kalman filter such as denoising and state estimation [53, 20]. The explicit inclusion of a (possibly nonlinear) Kalman filter becomes optional, thus lifting the requirement to identify the system equations. Further, RNNs have been used to describe actual neural dynamics and thus lend themselves to model sensorimotor impairment.

Other model choices may be desired to increase the biological realism using e.g. Wilson-Cowan equations [64], a spike-based neuron model [22], cortical microcircuits [2], hierarchical [1] or anatomical [33] constraints, or by explicitly including the resistive properties of neural tissue. To be compatible with our framework, the only requirement is that the model can be expressed in a form that supports automatic differentiation using e.g. Jax [8] or PyTorch [39].

Degradation or impairment of this neural system can be modelled by perturbing the RNN parameters. We demonstrated that its function can be restored by adding a secondary controller (modelling a prosthesis), which interacts with a subset of neurons in the neural system. We highlight three approaches to train the neural system and prosthesis, the choice of which depends on the properties and knowledge of the system.

### 4.2 Choosing the learning method

The three learning approaches considered here each come with their (dis-)advantages. Direct optimization of some cost function (Section 2.5.1) is the most efficient in terms of training data and time, but assumes a differentiable and known system, plus the existence of a suitable loss function. The use of an oracle teacher (Section 2.5.2) assumes a linear and known system, for which an analytic solution from optimal control theory can be computed. Then the training is similarly efficient as the direct method. Both achieve excellent accuracy as measured against the optimal control baseline and are a viable tool for the development of prosthetic controllers in silico. In practical settings, assumptions such as system knowledge or differentiability may not be satisfied, in which case one can resort to reinforcement learning. The outcome of RL-based training (Section 2.5.3) is less reliable and requires substantially more time, training data, and insight into relevant hyperparameters (though for the problems considered here, PPO defaults were sufficient). The major advantage of RL is its broad applicability even if the system equations are unknown, nonlinear, not differentiable, or only a part of the states is observed. Another promising feature of using RL is that it enables moving experimental design beyond mechanistic objectives (of neuronal activation) towards behavioral objectives (of reallife tasks) [27]. Finally, by using controllers based on deep neural networks, we gain access to powerful training tools from deep learning which are beneficial for neural control applications. In practice, the amount of training data will be limited, requiring sample-efficient [18, 26] or few-shot learning [60]. Plasticity in the neural system or a deterioration of the implant may require continuous adaptation of the prosthetic controller for which techniques from continual learning are readily available [38].

### 4.3 Extensions

The neural system RNN in the present work was trained to perform a control task in a simulated environment without constraining the resulting dynamics to approximate those observed in real neural systems. A next step is therefore to perform system identification using real neural recordings, similar to [41, 54]. Future work will also consider behavioral tasks in higher-dimensional environments, for instance with agents performing visual navigation in a real or virtual environment such as AI Habitat [56] or BEHAVIOR-1K [32]. Subsequent studies may also investigate different types of model degradation, e.g. simulating cell death or complete loss of sensory function.

We have shown in Section 3.3 that controllability and observability Gramians can be used to determine a lower bound on the number of stimulation and recording electrodes of an implant. The Gramians can be calculated analytically if the system is known and linear, but here we used a data-driven estimation technique [30] to demonstrate applicability to unknown nonlinear systems. In a behavioral setup, the required data could be obtained readily via psychophysical experiments. The Gramians indicate directions of efficient control and readout. A promising area of research concerns the combination of our work with the field of sparse sensor and actuator placement [34, 40, 11, 37], which will facilitate the optimal selection of stimulation and recording probes. This approach aligns well with recent studies that reveal stable low-dimensional latent dynamics of cortical neurons in a behavioral task [14].

Recent work [6] used Lyapunov theory to derive conditions for input-to-state stability in a multi-layer RNN. These conditions depend only on the learnable network parameters and can be used for safety verification, i.e., to ensure that outputs lie within a predefined safe region [25]. These results are applicable to the RNNs used here. Together with recent advances in safe RL [50], these techniques enable AI-based controllers that adhere to ethical standards [45, 21, 12] and pave the way to a translation into the clinical setting.

## 5 Conclusion

In light of these opportunities, we expect that the proposed framework will have a significant impact on the development of neural prostheses as it enables flexible in-silico testing of algorithms for stimulation and closed-loop control before validation using animal models or deployment in patients.

## Acknowledgements

This work has received funding from the European Union’s Horizon 2020 research and innovation programme under grant agreement No 899287.

## Footnotes

1 A black-box source of knowledge about a target function is sometimes called an ‘oracle’ in the machine learning community.

2 https://github.com/rbodo/neural_control

3 https://www.gymlibrary.dev/environments/mujoco/inverted_pendulum/

## Notes

### Competing Interest Statement

The authors have declared no competing interest.

